# BigDataProcessor2: A free and open-source Fiji plugin for inspection and processing of TB sized image data

**DOI:** 10.1101/2020.09.23.244095

**Authors:** Christian Tischer, Ashis Ravindran, Sabine Reither, Rainer Pepperkok, Nils Norlin

## Abstract

Modern bioimaging and related areas such as sensor technology has seen tremendous development the last years allowing several contemporary imaging techniques, electron microscopy (EM) and light sheet microscopy in particular, to generate datasets frequently reaching the size of several terabytes (TB). As a consequence, even seemingly simple data operations such as cropping, chromatic- and drift-corrections and even visualisation, poses challenges when applied to thousands of time points or tiles. To address this we developed BigDataProcessor2 – a Fiji plugin facilitating processing workflows for TB sized image datasets.

**Availability and implementation:** BigDataProcessor2 is available as a Fiji plugin via the BigDataProcessor update site. The application is implemented in Java and the code is publicly available on GitHub (https://github.com/bigdataprocessor/bigdataprocessor2).

## 1. Introduction

Inspection and processing of TB sized image data as produced by state-of-the-art light-sheet and volume electron microscopy poses several practical challenges (Power and Huisken, 2017). Even image inspection can be burdensome, because loading the entire dataset from disk into RAM, as it is usually done for conventional MB to GB sized data, is not feasible due to the limitations of a standard computer’s RAM. In addition, pixel wise image processing operations on the whole dataset can take hours and requires data duplication on disk as the processed images cannot be held in RAM.

The challenges of big image data inspection can be addressed by lazy-loading schemes where only the portion of the data is loaded into RAM that is needed to render the current view on the computer monitor. There are several commercial and open source solutions that adopt this strategy to interactively render big data. The rendering modes include fixed plane 2D slicing (Schneider et al., 2012), arbitrary plane 2D slicing and 3D volume rendering: Imaris (Oxford Instruments), Arivis Vision4D (Arivis AG), Amira (Thermo Fisher Scientific), BigDataViewer (Pietzsch *et al*., 2015), Vaa3D (Bria *et al*., 2015), TDat (Li *et al*., 2017). However, except for the fixed plane 2D slicing mode, these solutions require the data to be saved in specific blocked multi-resolution file formats that enable efficient loading of arbitrary (random access) data portions from disk into RAM and onto the GPU. Due to write performance considerations, raw microscopy data is typically not saved in a format that is compatible with those requirements. It is thus usually necessary to re-save the data in a specific file format that works with the respective image visualisation and analysis platform. Re-saving TB sized image data requires a significant amount of additional disk space and can take many hours and should thus ideally be only done once. In addition to the file format, raw microscopy data can have further issues. For example, only part of the acquired data may be of actual interest, either because larger fields of view have been acquired to compensate for unpredictable sample motion, or scientifically interesting phenomena have only occurred in specific parts of the imaged sample. Also, pixel density and bit-depth can be unnecessarily high, because camera-based microscope systems with fixed pixel size and bit-depth have been used. Moreover, there may be chromatic aberrations or other shifts between acquired data channels.

Taken together, to render raw microscopy data amenable for analysis it typically needs to be re-saved in a suited file format and a number of processing operations such as cropping, binning, bit-depth conversion, channel shift, and drift correction might need to be applied. It is important to realize that, if performed sequentially, each of these processing steps requires loading, processing, and re-saving of the entire (initially TB sized) dataset, which, taken together, could take several hours or even days. It would be much more efficient to load the raw data, apply all processing steps in RAM and then re-save the data only once.

To this end, we developed BigDataProcessor2 to facilitate interactive browsing and initial processing of microscopy raw data, before, in one go, re-saving the processed data, ready for image visualisation and further analysis.

## 2. Implementation and Application

Our first implementation (BigDataProcessor1) was written in Java, making use of ImageJ’s VirtualStack class (Schneider et al., 2012) for lazy-loading of big image data. Here, we focus on our new implementation (BigDataProcessor2, in short BDP2), which is built on a more recent Java library for image processing, namely ImgLib2 (Pietzsch et al., 2012). The source code for both implementations is publicly available on GitHub (https://github.com/bigdataprocessor) and the applications can be installed via a Fiji (Schindelin et al., 2012) update site “BigDataProcessor”. All functionality is accessible to users without the need for programming via a graphical user interface (GUI).

Currently, BDP2 supports inspection and processing of 5-D (x, y, z, channel and time) data from collections of single- and multi-plane Tiff as well as HDF5 (The HDF Group, 1997) based file formats, thereby accommodating a majority of raw data formats currently occurring in light sheet and volume electron microscopy (see Supplementary Information). The loading can be freely configured using regular expressions and, for convenience, some frequently occurring loading schemes are preconfigured already. BDP2 employs Imglib2’s `CachedCellImg` class for lazy-loading of image data from files into memory. The `CachedCellImg` partitions an image into small blocks (cells), where a cell’s content is read from disk only when the application requests access to this part of the image. For image visualisation, BDP2 employs BigDataViewer, BDV in short (Pietzsch et al., 2015), which provides arbitrary plane slicing of volumetric, multi-channel time-lapse image data. As one microscopy camera image typically contains only a few MB of data, it fits readily into RAM and can be loaded within (sub-)seconds, given a data access bandwidth larger than or equal to a few MB/s. These bandwidths are nowadays typically available and as such BDP2’s lazy-loading scheme allows for interactive browsing of TB sized image data on a standard computer.

**Figure 1.**
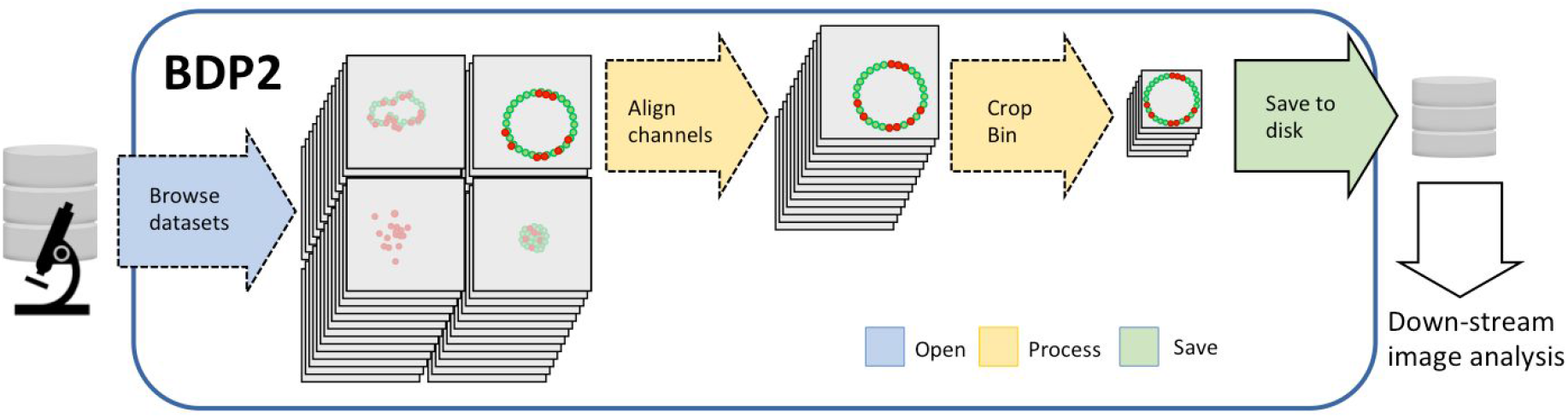
Schematic representation of a lazy-processing workflow in BigDataProcessor2 (BDP2). Dashed arrows represent lazy-computation, where only the pixels needed to render the currently viewed image plane are loaded and processed. The complete data browsing, data selection and data processing workflow can be configured in a few minutes even for TB-sized image data. Only the final saving to disk requires processing of the whole dataset and will take a correspondingly long time (up to hours).

Importantly, all image processing operations are performed lazily, using Imglib2’s `Views` and `Converter` classes. This limits computations to the pixels needed to render the currently viewed image plane. At present we support the following lazy processing operations: affine transformed viewing, cropping, binning, bit-depth conversion, and channel alignment (e.g., chromatic shift correction, split chip acquisition). The compute times for these operations are in the sub-second range such that the user can interactively configure all image processing steps while inspecting arbitrary locations in the sample (see Supplementary Figure 1 and Supplementary Movie 1,2). All operations are accessible via a GUI, but can also be recorded and executed as ImageJ macro scripts making it possible to readily share and publish the processing workflow. A detailed description of all available menu items can be found in the Supplementary Information. Once all processing steps have been configured, the dataset can be re-saved in a format suitable for further analysis. We currently provide saving in open-source Tiff and Hdf5 based formats, where we save one file for each 3D volume. For TIFF, we also support saving each plane to a separate file. This is useful, e.g., for EM data where already a single 3D volume can be several TB. For HDF5, we save the data in a chunked pyramidal format allowing for efficient viewing with both BDV and Imaris (Imaris v9.0, Bitplane AG). In all cases we support lossless compression algorithms (see Supplementary Information).

**Figure 2.**
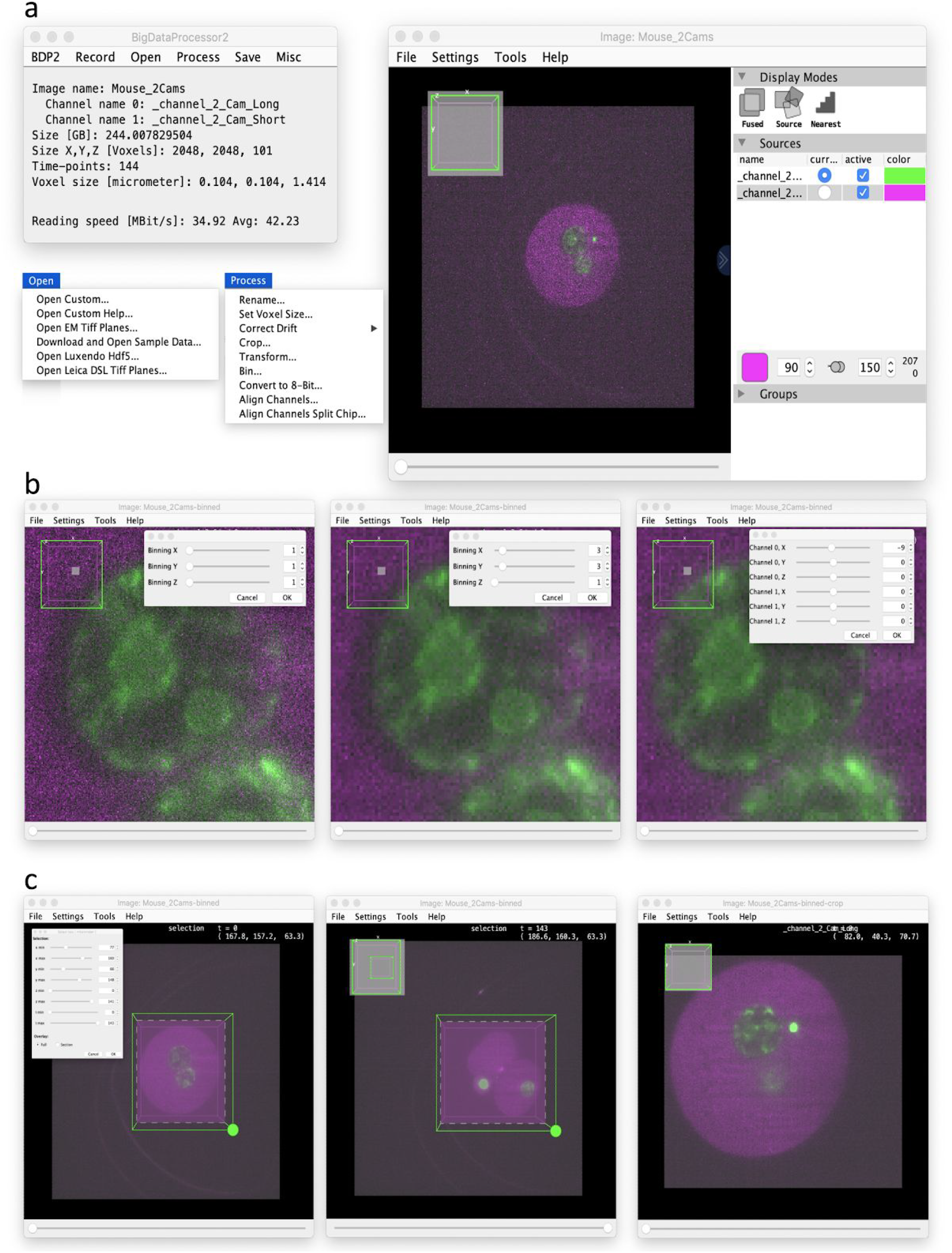
Overview of the user interface and an example workflow in BigDataProcessor2 (BDP2). (**a**) Image browsing. Left: The main BDP2 window with the Open and Process dropdown menus expanded. Right: BigDataViewer’s user interface. (**b**) Binning and channel alignment. Left: Original data (zoomed in). Middle: 3 x 3 binning in X & Y, reducing noise, and the size of the dataset by a factor of 3 x 3 = 9. As all processing operations, also the binning is performed by means of lazy computation and can thus configured interactively even for TB sized datasets. Right: Channel alignment, correcting a shift between the green and magenta channel (note the difference to the middle panel). (**c**) Cropping. Left: First time point of the dataset with interactive cropping user interface. Middle: Last timepoint of the dataset with enlarged cropping area to include all relevant data. Right: Cropped data, showing again the first timepoint. Thanks to BDP2’s lazy-loading and lazy-processing the above steps (a-c) can be executed in a few minutes. Finally, the data could be re-saved using the Save menu (not shown). In this example, binning and cropping helped to reduce the size of the data from 244 GB to 4.8 GB without loss of biologically relevant information.

## 3. Discussion and Conclusions

Data bandwidth limitations are an issue of rising importance designing modern bioimage analysis pipelines, especially with increasing computer virtualisation and non-local data storage. Therefore, the benefits of optimised data transfer to only deliver the data in use and minimise excessive reading from files or parts of files cannot be underestimated. Recognising this current bottleneck sparked the work on BigDataProcessor2. Even at a bandwidth of 100 MB / s loading just a typical sized single time-point light sheet volume (~1 GB) from disk into RAM would take ~10 s, which is impractical for data inspection. However, at the same bandwidth a 4 megapixel 16-bit image plane can be loaded in less than 1 s. Thus, a plane-wise lazy-loading scheme (such as implemented in BigDataProcessor2) allows for interactive browsing through a TB sized microscopy raw dataset. Importantly, using Imglib2, BDP2 also provides lazy processing steps that do not noticeably increase the latency, because computations are only performed on pixels needed to render the currently viewed image plane. Thereby data can be inspected and a processing pipeline can be interactively configured within a few minutes. However, the actual processing and re-saving of a large dataset can take up to several hours. This is generally not a problem in practice, since all processing steps have been configured and no more human interaction is required during the saving. In our experience resaving has most often resulted in a significant (up to 100 fold) data size reduction and has allowed us to tape backup the raw data in favour of a resaved processed version of the data.

In conclusion, we consider the BigDataProcessor2 Fiji plugin to significantly simplify the inspection and processing of big image data. As big image data is becoming increasingly prevalent we are positive that the already existing user base (e.g. Alladin et al. 2020, Villani et al. 2019, Wolny et al. 2020) will grow even further in the future.

## Acknowledgements

We thank the EMBL IT-department for support, Tobias Pietzsch (MPI CBG, Dresden) for help related to BigDataViewer, Ashna Alladin, Matthew Boucher, Isabell Schneider (EMBL Heidelberg), Gustavo Quintas Glasner de Medeiros (FMI Basel), Alexis Maizel (University Heidelberg), Marion Louveaux (Institut Pasteur), Jan Rhoden (Bruker), Bjoern Eismann (Bruker) and the Francesca Peri lab (University of Zürich), for testing, reading the manuscript and/or providing suggestions.

## Supplementary Information

### Supplementary Note 1: Menu items

**Supplementary Figure 1:**
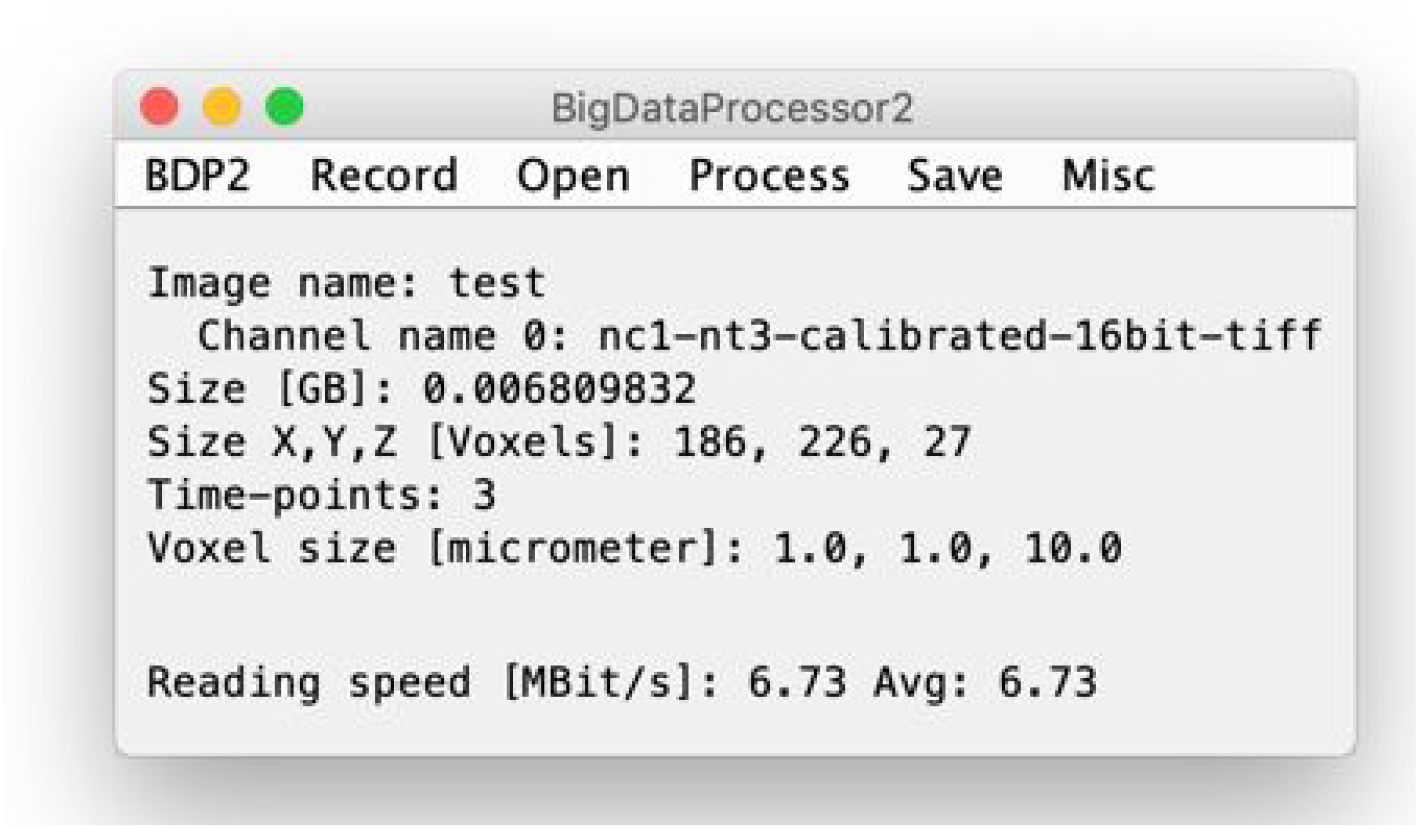
Screenshot of the main user interface.

BDP2 comes with its own user interface (UI) where all functionality can be accessed (the menus in the ImageJ UI will typically not work here). The UI shows information about the currently active image as well as the current and average image data reading speed. It is possible to have multiple images (BigDataViewer windows) open at the same time. Following the usual ImageJ convention, the “active” image is the one that you clicked on last.

**Supplementary Figure 2:**
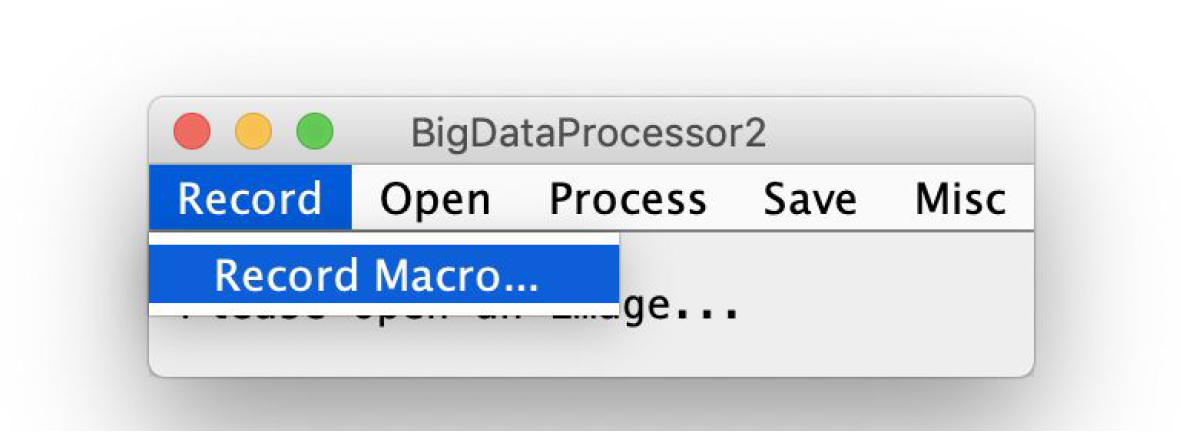
Screenshot of the Record menu.

#### Record > Record Macro…

Enable/ disable macro recording. This is equivalent to [Plugins > Macros > Record] in the ImageJ menu.

##### Motivation

Macro recording is one of ImageJ’s greatest features as it allows users without programming experience to record reusable scripts. It can be used for automation but also for documentation.

**Supplementary Figure 3:**
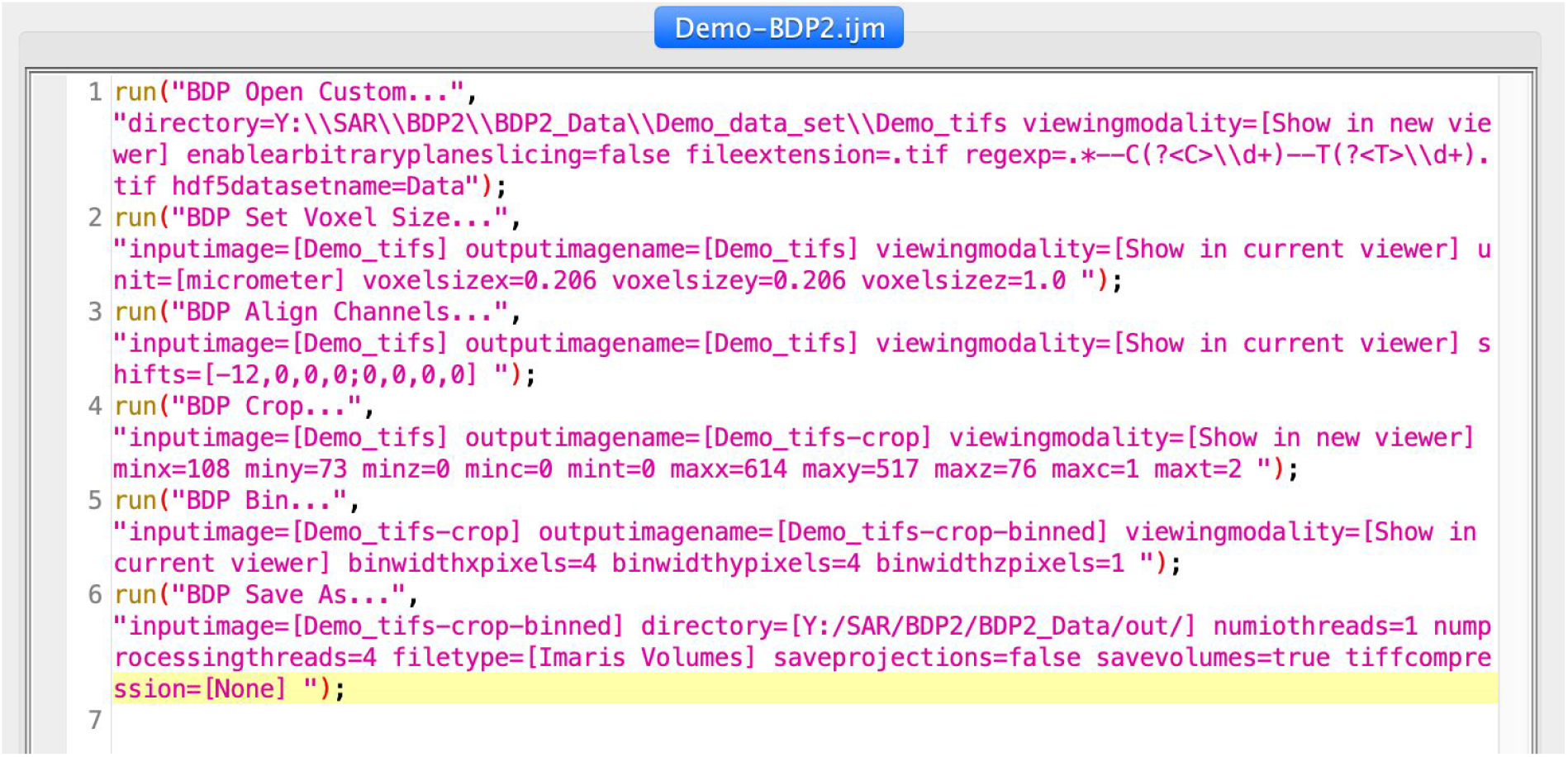
Screenshot of a recorded macro.

**Supplementary Figure 4:**
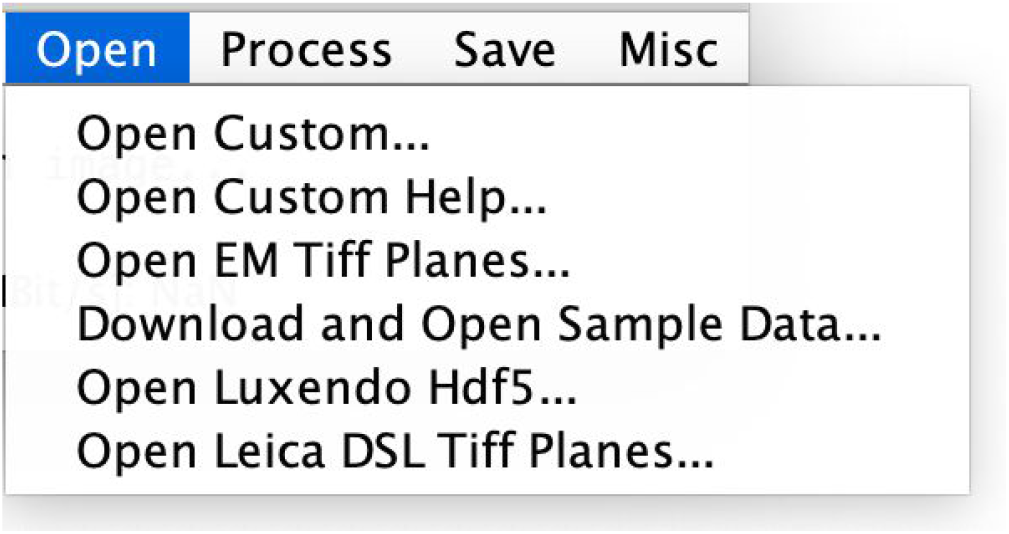
Screenshot of the Open menu.

### Open > Open Custom…

Open datasets consisting of a collection of Tiff or Hdf5 volumes. The assignment of each file (volume) to a channel and time point can be specified by a regular expression.

### Open > Open Custom Help…

Shows and explains a number of regular expressions that can be used in the [Open > Open Custom…] menu item.

### Open > Open EM Tiff Planes…

Opens a single folder with Tiff single plane files. Each file will be assigned to one z-plane in a dataset with one color and one time point.

#### Motivation

This is a typical format for volume EM data to be stored in.

### Open > Download and Open Sample Data…

Download and open sample data stored in the BioStudies archive (https://www.ebi.ac.uk/biostudies/studies/S-BSST417?query=bigdataprocessor2).

#### Motivation

Conveniently accessible example data is useful to explore/teach BigDataProcessor2 without the need to prepare suitable input data.

### Open > Open Luxendo Hdf5…

Open datasets acquired with Luxendo light sheet microscopes.

#### Motivation

Luxendo uses an open-source hdf5 based file format. We added convenience functionality for opening those files without the need to enter a complex regular expression.

### Open > Open Leica DSL Tiff Planes…

Open datasets acquired with Leica DSL microscopes, choosing “Auto-Save, Data type: Tif, Compression: Uncompressed” as an option (Leica’s proprietary file format is called .lif, which we do not currently support).

#### Motivation

While the Tiff file format is open source, Leica’s naming scheme would require entering a complex regular expression and we thus implemented this convenience opening functionality.

**Supplementary Figure 5:**
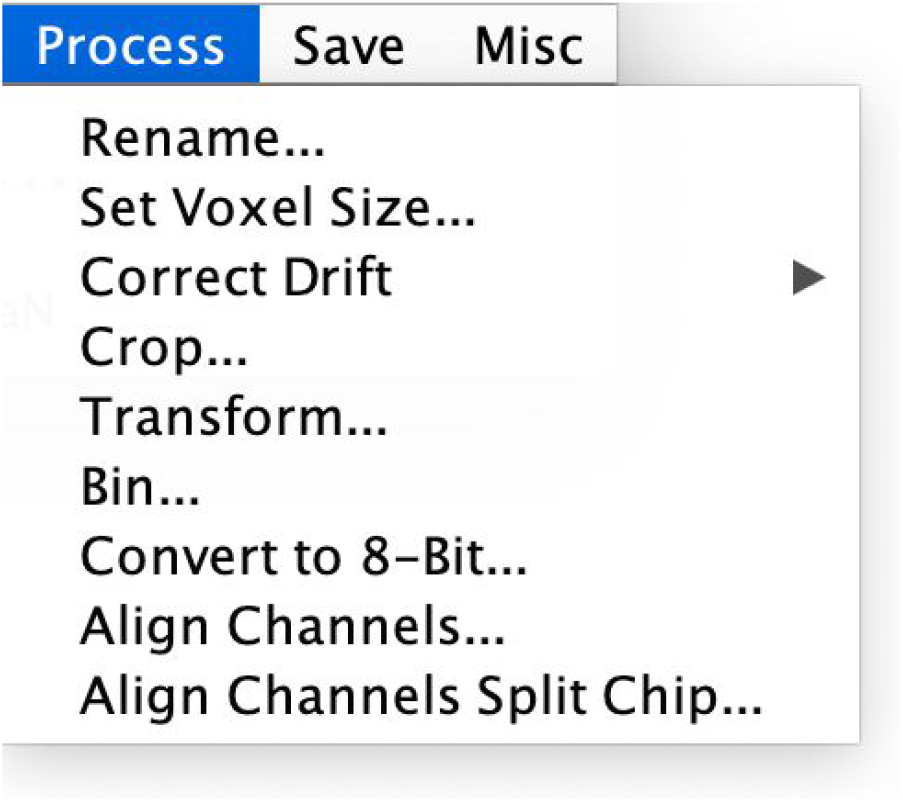
Screenshot of the Process menu.

### Process > Rename…

Rename the dataset.

### Process > Set Voxel Size…

Change the voxel size image properties.

#### Motivation

The voxel size may not always be read correctly from the dataset, thus it is useful to have the option to set it manually.

### Process > Correct Drift

Correct sample motion by interactively creating a 3D track, which will be applied such that the image is stationary relative to the track positions.

#### Motivation

For time lapse data there is a risk that a sample moves during acquisition. To accommodate for either sample or microscope drift it is common to choose a field of view to encompass expected drift at the expense of larger data footprint. This can be compensated by cropping the data. However, applying a static volumetric crop over the whole time lapse is suboptimal. Therefore an ideal crop would be on drift corrected data (see Supplementary Movie 2). Additional applications can be, e.g., tracking motile cells in tissues.

### Process > Correct Drift > Create Track…

Create a 3D track by manually placing anchor points in a subset of time points (track positions in the other time-points will be automatically added by linear interpolation). When done, save the track as a Json file to disk, to be used in [Process > Correct Drift > Apply Track…].

### Process > Correct Drift > Apply Track…

Load a 3D track from a file (created with [Process > Correct Drift > Create Track…]) and apply it to the dataset. This will cause the dataset to be displayed with each timepoint shifted according to the track positions (no data duplication).

### Process > Crop…

Interactively specify a 4D (x,y,z,t) subset of the data to be displayed in a new viewer window.

#### Motivation

Imaging processes in living samples require setting up imaging parameters before knowing exactly when and where the process of interest takes place. Therefore the imaging field of view (x,y,z) and temporal extent (t) are usually set generously to accommodate sample drift, motion, or growth. Using the crop function one can reduce the dataset to the necessary spatial and temporal dimensions.

### Process > Bin…

Performs arbitrary binning along x y and z coordinates.

#### Motivation

For camera-based microscopy systems the effective pixel size often cannot be freely chosen during acquisition. Thus, the user may be forced to oversample, leading to large data volumes and potentially significantly increased image processing times. Thus, binning the data post-acquisition can be very useful as it both reduces data size (and noise), often without compromising scientific accuracy.

### Process > Convert to 8-bit…

Convert the dataset from 16 to 8-bit depth.

#### Motivation

Cameras typically produce image data at 12, 14, or 16 bit-depths. For many image analysis tasks, 8-bit depth is sufficient affording the user to reduce data size by a factor of 2. However, converting 16-bit to 8-bit data is not trivial as it entails deciding on a specific mapping from the higher to the lower bit-depth, which will lose information. Choosing a mapping of 65535 to 255 and 0 to 0 can lead to a low dynamic range in the 8-bit range especially when the input contains only a subset of the full 16-bit range. Also mapping max(image) to 255 and min(image) to 0 can be sub-optimal if there are spurious pixels with very high values, again leading to a low dynamic range for the relevant grey values in the 8-bit converted data. We thus provide the possibility to interactively specify a mapping while browsing the dataset to inspect the result of the applied conversion.

### Process > Align Channels…

Shift one channel in relation to the other to compensate pixel offsets e.g. due to chromatic shifts.

#### Motivation

Chromatic shifts either due to optics being corrected only for a given wavelength range, or parallel acquisition of two channels on two cameras can lead to offsets between the two channels/ images. We, therefore, provide the functionality to correct for such channel shifts in x,y and z.

### Process > Align Channels Split Chip…

Specify two crop regions in one channel and convert those regions into two channels, i.e. the number of channels of the resulting image is increased by one.

#### Motivation

For the sake of acquisition speed, some fluorescence microscope systems acquire the signal of several fluorescence channels simultaneously on the same camera chip. Thus, we provide the functionality to convert such data into a conventional multi-channel dataset by aligning the channels from a “split chip”.

### Process > Transform…

Renders an affine view of the data.

#### Motivation

Useful when data is warped due to an acquisition process that renders x-y-z non-orthogonal. Examples are when a stage movement is not orthogonal to the field of view. Also useful in single objective light sheet microscopy.

**Supplementary Figure 6:**
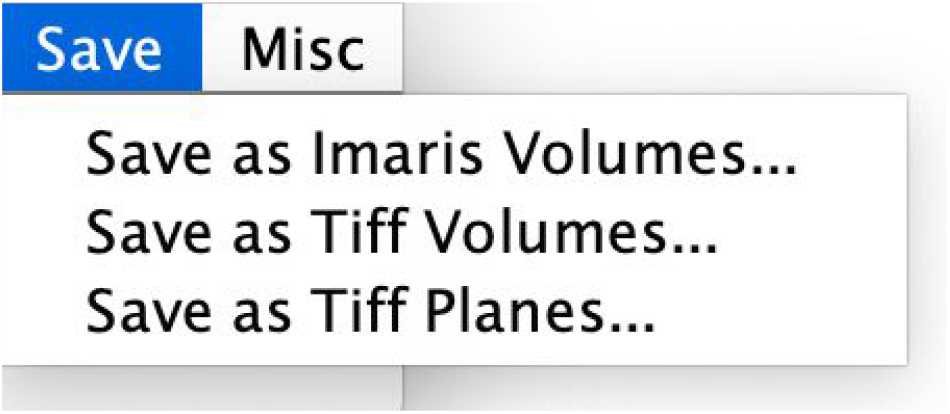
Screenshot of the Save menu.

### Save > Save as Imaris Volumes…

Save dataset as an hdf5 based pyramidal Imaris file (http://open.bitplane.com/ims), with each channel and time point saved as an individual .h5 file and one .ims header file that can be used to view the data both in Fiji’s BigDataViewer and in the commercial Imaris software.

#### Motivation

The low data overhead of a pyramidal scheme (in 3D for binning 2 x 2 x 2 at each pyramidal level ~14%) is a marginal cost for a substantially improved user experience when viewing the data. We, therefore, provide saving data in an open file format that offers this functionality based on hdf5, which means that it can be handled with all common programming languages.

### Save > Save as Tiff Volumes…

Save the dataset as a series of Tiff stacks with each channel and time point saved as an individual .tif file.

#### Motivation

Tiff stacks are still the most used and compatible file format that can be easily opened by all software for downstream analysis.

### Save > Save as Tiff Planes…

Save the dataset as a series of Tiff planes, where each z-slice, channel and time point are saved as an individual .tif file.

#### Motivation

Saving a volume as a series of Tiff planes is popular e.g. in the EM community.

**Supplementary Figure 7:**
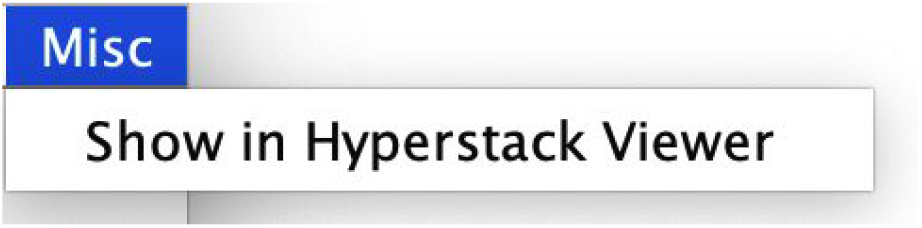
Screenshot of the Misc menu.

### Misc > Show in Hyperstack Viewer

Opens the current image virtually in the “classic” ImageJ hyperstack viewer.

#### Motivation

BigDataViewer is a relatively recent addition to the ImageJ ecosystem and many users like to use the classical ImageJ hyperstack viewer. In addition, with the data being displayed in the hyperstack viewer, one has access to many useful inspection tools such as intensity histograms and intensity line profiles.

### Supplementary Movie 1: BigDataProcessor2 Workflow

https://www.youtube.com/embed/OixZ0ILbkvc?vq=hd1440

#### Opening, browsing and preprocessing of a 250GB Hdf5 based raw dataset

Movie shows a screen recording of a basic preprocessing workflow of a 250 GB Hdf5 image dataset acquired by light-sheet microscopy. The following steps are demonstrated: Open Luxendo HDF5 -> Brightness & Color adjustment -> Set Voxel Size -> Align Channels -> Crop -> Bin -> Save. The 2 color early mouse embryo data were provided by Manuel Eguren, Ellenberg group EMBL Heidelberg.

### Supplementary Movie 2: Drift Correction

https://www.youtube.com/embed/7SCZlToxY9E?vq=hd1440

#### Drift correction of a 250 GB HDF5 dataset

The movie shows a screen recording of the drift correction of a 250 GB Hdf5 image dataset acquired by light-sheet microscopy. The following steps are shown in the movie: Correct Drift -> Create Track -> Apply Track. The single color mouse mammary gland organoid data were provided by Ashna Alladin, Jechlinger group, EMBL Heidelberg.

## References

Alladin A., Chaible L., Garcia del Valle, L., Reither S., Loeschinger M., Wachsmuth M., Hériché J.K., Tischer C. and Jechlinger M. (2020). Tracking cells in epithelial acini by light sheet microscopy reveals proximity effects in breast cancer initiation. Elife 2020;9:e54066

Bria A., Iannello G., Onofri L. and Peng H. (2016) TeraFly: real-time three-dimensional visualization and annotation of terabytes of multidimensional volumetric images. Nature Methods, 13, 192–194.

Imaris v9.0 Bitplane AG.

Li Y., Gong H., Yang X., Yuan J., Jiang T., Li X., Sun Q., Zhu D., Wang Z., Luo Q. and Li A. (2017) TDat: An Efficient Platform for Processing Petabyte-Scale Whole-Brain Volumetric Images. Front. Neural Circuits, 11, 51.

Pietzsch T., Saalfeld S., Preibisch S. and Tomancak P. (2015). BigDataViewer: visualization and processing for large image data sets. Nature Methods, 12, 481–483.

Pietzsch T., Preibisch S., Tomancak P. and Saalfeld S. (2012). ImgLib2 – generic image processing in Java. Bioinformatics, 28, 3009–3011.

Power R.M. and Huisken, J. (2017) A guide to light-sheet fluorescence microscopy for multiscale imaging. Nat. Methods, 14, 360–373.

Schindelin J., Arganda-Carreras I., Frise E., Kaynig V., Longair M., Pietzsch T., Preibisch S., Rueden C., Saalfeld S., Schmid B., Tinevez JY., James White D., Hartenstein V., Eliceiri K., Tomancak P. and Cardona A. (2012) Fiji: an open-source platform for biological-image analysis. Nat. Methods, 9, 676–682.

Schneider C., Rasband W. and Eliceiri K. (2012) NIH Image to ImageJ: 25 years of image analysis. Nat. Methods, 9, 671–675.

The HDF Group (1997) The HDF Group. Hierarchical Data Format, version 5, 1997-NNNN.

Villani A., Benjaminsen J., Moritz C., Henke K., Hartmann J., Norlin N., Richter K., Schieber N., Franke T., Schwab Y. and Peri F. (2019) Neuronal clearance by microglia depends on packaging of phagosomes into a unique cellular compartment. Dev. Cell, 49 77–88

Wolny A., Cerrone L., Vijayan A., Tofanelli R., Vilches Barro A., Louveaux M., Wenzl C., Steigleder S., Pape C., Bailoni A., Duran-Nebreda S., Bassel G., Lohmann J., Hamprecht F., Schneitz K., Maizel A. and Kreshuk A. (2020) Accurate and Versatile 3D Segmentation of Plant Tissues at Cellular Resolution. bioRxiv 10.1101/2020.01.17.910562v1

